# The Insertion Sequence Excision Enhancer (IEE): a PrimPol-based system for Immobilizing Transposon-Transmitted Antibiotic Resistance Genes?

**DOI:** 10.1101/2023.06.07.543989

**Authors:** Mick Chandler, Karen Ross, Alessandro M. Varani

## Abstract

We provide an overview of a protein, IEE (Insertion Sequence Excision Enhancer), which was originally observed to facilitate high levels of excision of the IS3 family member, IS629, from clinically important Escherichia coli O157:H7. IEE was subsequently shown to affect a large class of bacterial insertion sequences which all transpose by producing a circular intermediate and presumably use the copy-out-paste-in transposition mechanism. Excision is dependent on both IEE and transposase indicating that it is associated with the transposition process itself. We propose that IEE serves to immobilize genes carried by compound transposons by removing the flanking IS copies, an activity which would explain the presence of certain of these genes without associated IS copies in plasmids and chromosomes. The biochemical activities of IEE as a primase with the capacity to recognize microhomologies, together with the observation that its effect appears to be restricted to those IS families which probably use the copy-out-paste-in transposition pathway, suggests a strand switch mechanism during the copy-out step leading to abortive transposition. This reinforces the proposal made for understanding the loss of the IS*30* family member, IS*Apl1*, which flanks the *mcr*-1 gene in the compound transposon Tn*6330*.

## Introduction

Bacterial insertion sequences, short segments of DNA capable of movement from one DNA site to another, are an abundant component of bacterial genomes and play important roles in the spread of host genes and often in the expression of these genes. Often, two (nearly) identical IS copies are found flanking specific genes in both plasmids and chromosomes and the entire structures are capable of transposition. These are known as compound or composite transposons (see (1)). Many, but certainly not all, such transposons carry antibiotic resistance passenger genes and are an important source of such genes.

Insertion sequences have been classified into nearly 30 different families (ISfinder; (2)) based on various characteristics which include the sequence of their transposases, their ends (IR) and the number of base pairs duplicated at the target site associated with insertion (DR) (3). Although the majority of IS encode a transposase with a DDE RNaseH fold, indicating the triad of amino acids at the catalytic heart of the enzyme, different IS families use different transposition pathways. Generally, DDE transposases appear to catalyze 3’ cleavage at each of the transposon tips, producing a 3’OH. (The cleaved strand is known as the transferred strand because it is “transferred’ into the target site). They process the 5’ cleavage in different ways (4,5). Two major pathways for the subsequent steps are known. One is called “cut-and-paste” where the free 3’OH generated by the first cleavage is used as a nucleophile to attack the second strand, shedding the flanking DNA and forming a small hairpin structure at the transposon ends. This then undergoes a second cleavage to generate the final transposition intermediate. In the second principal pathway, known as “copy-out-paste-in” (6), the 3’OH generated by the first cleavage is used as a nucleophile to attack the same strand at the opposite transposon end. This generates a single strand bridge and a second 3’OH which acts as a primer for synthesis of a circular transposon copy (see below for details) with abutted left and right ends. The abutted ends are highly active in transposition and the circle undergoes transposase-mediated integration into its chosen target site.

Although only a few have been analyzed in any detail, a large number of the IS families including IS*1* (7,8), IS*3* (9,10), IS*21* (see(11)), IS*30* (12), IS*256* (13,14), IS*110*/IS*1111* (see(15–19)), IS*Lre2* (20), IS*L3* (21,22) and IS*1202* (23,24) have circular transposition intermediates and presumably transpose by a copy-out-paste-in mechanism.

Evolutionary studies often provide evidence that genes have been shared between different organisms based on their DNA and protein sequence relationships and the phrase Lateral or Horizontal Gene Transfer is often used in this context. However, they rarely provide any mechanistic information concerning how these gene transfers might have occurred. With the astonishing increase in the number of sequences in the public databases over the past two or three decades, identical antibiotic resistance genes have been identified in different sequence contexts with and without flanking IS. Major outstanding questions include how these passenger genes have been assimilated into compound transposons and whether the immobile, IS-less, genes have been derived from active compound transposons by deletion of the flanking IS.

Studies from the Oka lab several years ago identified a gene, called <u>Insertion Sequence Excision enhancer or IEE</u>, whose product appeared to stimulate IS excision by several orders of magnitude (25). This gene product might stimulate excision of the flanking IS without leaving any discernable scars and thus provide a potential explanation for the extended presence of certain genes, normally carried as part of compound transposons, in contexts without associated IS.

Below we summarize present knowledge concerning IEE, and provide a model based on our studies on the *mcr* colistin resistance gene and its associated compound transposon Tn*6330* (26–29) for its ability to immobilize transferred genes while leaving only the most subtle scars in the process.

### IS Excision is Stimulated by High Transposase Levels

The IS*3* family insertion sequence IS*1203*v (similar to IS*629*) was identified in a Shiga toxin 2 gene (*stx2*) of *Escherichia coli* O157:H7, which it had insertionally inactivated. When supplied with high levels of transposase, the IS underwent precise excision leading to reactivation of *stx2* (30).

An assay, which measured reversion to ampicillin resistance from a single copy F plasmid derivative in which an ampicillin resistance gene was interrupted by an IS*1203*v insertion, *bla*::IS*1203*v, was used to monitor precise excision rates. When the *bla*::IS*1203*v-carrying plasmid (Fig.1A) vector was introduced into both *E. coli* O157:H7 and *E. coli* K12(MG1655), excision was observed to occur approximately 10^5^ fold less frequently in *E. coli* K12 than in the *E. coli* O157:H7 host (25). It was noted that *E. coli* O157:H7 was known to carry a significant number of IS*629* copies whereas *E. coli* K12 was devoid of this IS (31,32), and it was thought that these might contribute in some way to the excision process. Further studies using a number of *E. coli* isolates with and without IS*1203*v/IS*629* copies supported the idea that excision was higher in those strains already carrying the IS.

Like other members of the IS*3* family, IS*1203*v (Fig. 1B) encodes an upstream gene, *orfA*, which presumably carries sequence-specific DNA binding functions, and a downstream reading frame, *orfB*, which is arranged in the -1 reading phase compared to *orfA* and includes the catalytic triad DDE signature. Programmed -1 translational frameshifting (33–35) between of *orfA* and *orfB* produces a gene product *orfAB* which functions as the transposase (see (6,36)). In an experimental system in which various transposition functions were supplied *in trans* from a compatible plasmid (Fig. 1C), excision was observed to be very low or below the detection level in the MG1655 host. In the O157:H7 strain, however, supplying the OrfAB transposase induced high deletion levels (⋃10^−3^) compared to those obtained with the empty vector (7.8x10^−7^) whereas supplying the *orfA, orfB* and *orfAB* genes in their native configuration only resulted in a moderate frequency of excision (2.6x10^−6^) and supplying *orfA* alone depressed excision (3.8x10^−9^). This implies that it is the level of available OrfAB transposase that is a determining factor in excision (25).

**Figure.1.**
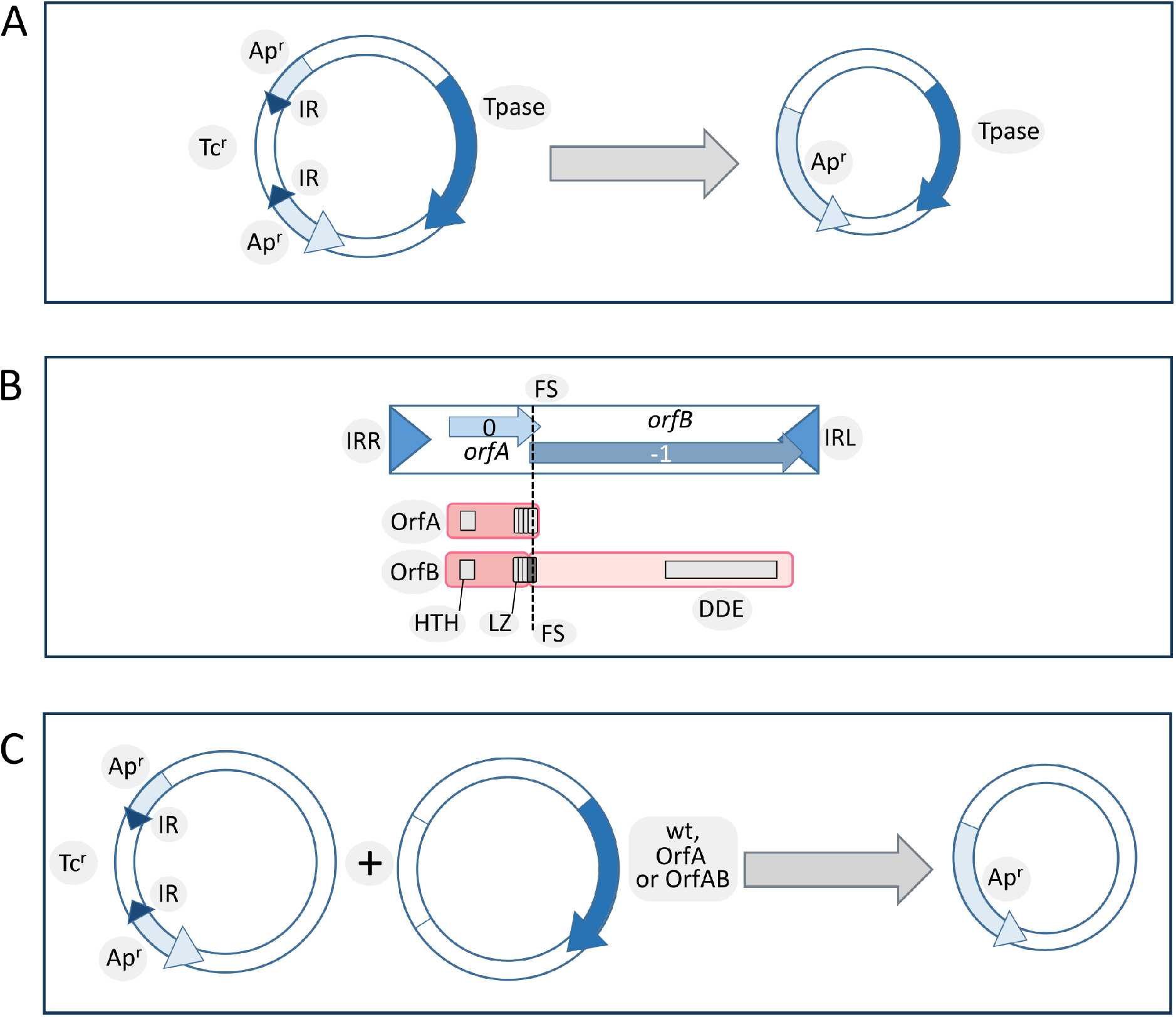
(A) System used to monitor IS excision. Redrawn and adapted from Kusumoto et al 2011 PMID: 21224843. The test plasmid (left) carried an insertion in an ampicillin resistance gene (bla::IS*1203*v/IS*629*, light blue) of a disabled IS composed of two terminal inverted repeats (IR, dark blue triangles) flanking a tetracycline resistance gene carried by a plasmid containing an orfAB transposase gene (dark blue arrow). Excision reulsts in loss of tetracycline reistance and appearance of ampicillin resistance (right) **(B) Genetic organization of IS*3* family members**. The IS is shown as a box. The triangles at each end represent the left (IRL) and right (IRR) terminal inverted repeats. The two open reading frames, orfA (light blue) and orfB (dark blue) are positioned in relative reading phases 0 and −1, respectively, as indicated. The region of overlap between *orfA* and *orfB*, includes the frameshifting signals to produce OrfAB,. The point at which the frameshift occurs, within the last heptad of the LZ, is indicated by the vertical dotted line. Below is a structure function map of OrfAB and OrfA. HTH, a potential helix-turnhelix motif; LZ, a leucine zipper motif involved in homo- and hetero-multimerization of OrfAB and OrfA. The catalytic core of the enzyme which the DDE triad. **(C) System used to test different configurations of IS*1203v*/IS*629* transposition genes**. This is composed of the bla::IS*1203*v/IS*629* cassette carried by onle plasmid and the transposition genes carried by a second, compatible plasmid.

### A Dedicated Enzyme: Identification of a common reading frame ECs1305 in all high excision strains

The authors also identified a reading frame, ECs1305, present in all high excision strains that was absent in the low excision strains (37). It was identified both in enterohemorrhagic (EHEC) and enterotoxigenic (ETEC) *E. coli* strains, and homologues were also identified by Blast analysis in a broad range of bacteria including Alpha-Beta-, Gamma-, Delta- and Epsilon-proteobacteria; *Bacteroides*; *Chlorobi*; *Cyanobacteria*; *Firmicutes*; *Actinobacteria*; and *Verrucomicrobia* (37).

More recently it has been estimated that a highly conserved gene copy is present in over 30% of available *E. coli* genome assemblies and is very abundant not only within enterohemorrhagic and enterotoxigenic genomes but also within enteropathogenic *E. coli* (38). The ECs1305 gene was subsequently named ***iee*** for **I**S-**e**xcision **e**nhancer (37). In EHEC O157, ECs1305/IEE is located in a large potential integrative element that is similar to SpLE1 (39) and has probably been dispersed in this way.

### IEE and an active transposase are required for high level IS excision

When the *iee* reading frame was deleted, the IS excision frequency measured by the bla::IS*1203*v/IS629 reversion as a readout, was greatly decreased but could be restored by reintroduction of a plasmid-carried *iee* copy. Moreover, the use of DDE mutant transposases was used to demonstrate that the ECs1305-promoted excision behavior was also dependent on an active transposase (37).

The effect on excision frequency of a number of host genes involved in DNA repair, structuring or processing, was investigated and although some of these had been shown to affect excision of other ISs (e.g.(40–42), the effects of mutation in these genes were largely marginal (Fig.2) and not as pronounced as mutations in ECs1305/iee (37).

**Figure.2.**
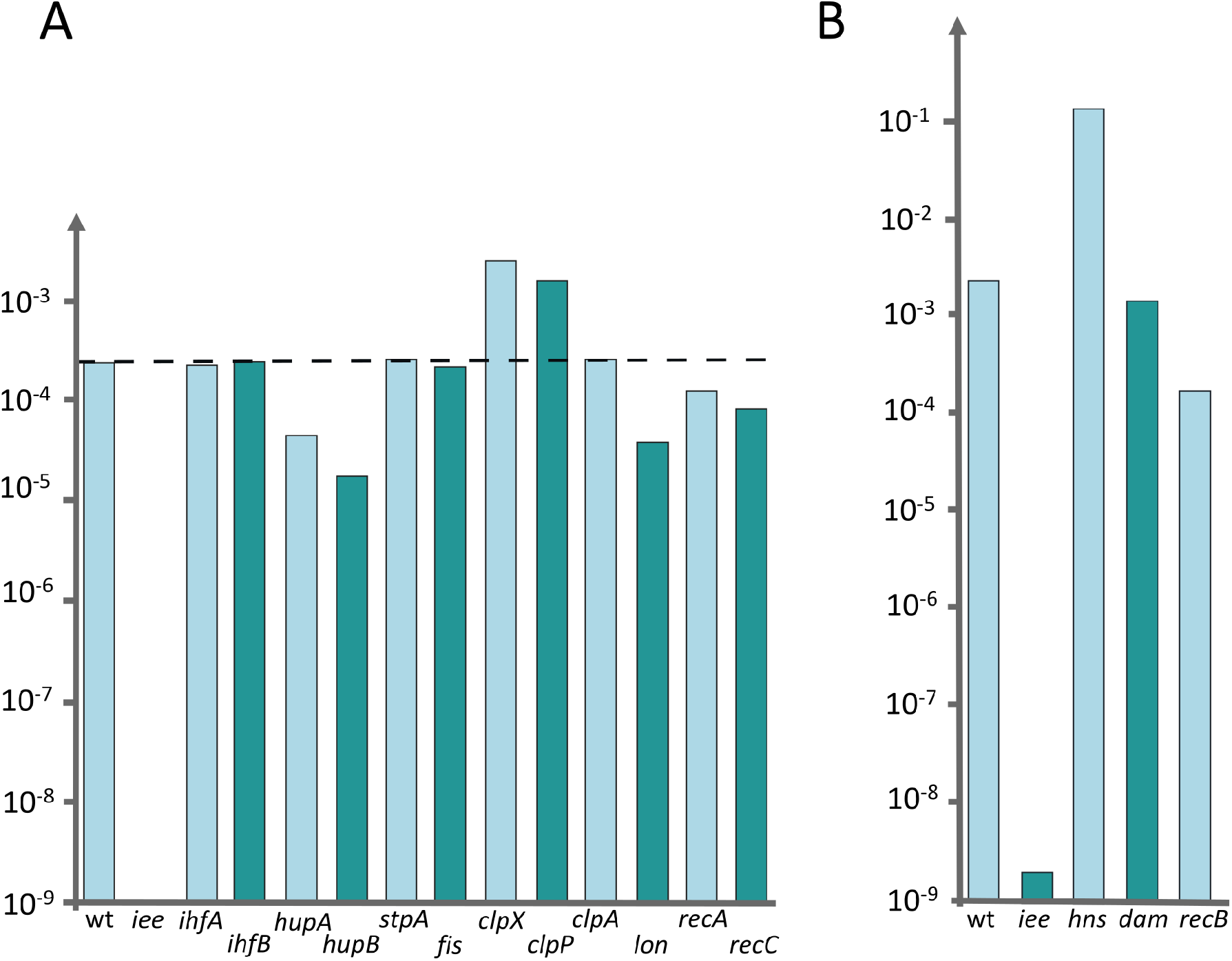
A Histogram showing the frequency of excision of IS*629* in various Mutant *E. coli* Genetic Backgrounds. Redrawn and adapted from Kusumoto et al 2011 PMID: 21224843. The assay consisted of a plasmid carrying an ampicillin resistance (Ap^r^) gene inactivated by insertion of a copy of IS*629* whose transposase had been substituted for a tetracycline resistance gene (Tc^r^). The plasmid also included the transposase gene placed under control of an external promoter. Excision of the IS*629* derivative results in loss of Tc^r^ and appearance of Ap^r^. The authors used a number of deletion mutants of genes which are thought to influence various aspects of transposition: IHF (integration host factor), HU, H-NS, FIS (factor for inversion stimulation), ClpXP5 protease Lon protease, Dam, RecA, and RecBC. The precise IS*629* excision frequency was examined in each mutant using the reporter plasmid-based assay. The hns, dam, and *recB* deletion mutants could not be generated in the original *E. coli* O157 Sakai strain, and were generated in another *E. coli* host carrying a chromosomally inserted *iee* gene.

### Analysis of the primary Sequence of IEE: a PrimPol Helicase

Alignment of a number of IEE proteins revealed 4 conserved regions (37) (Fig.3). Two of the more N-terminal regions (1 and 2) showed similarities to eukaryotic/archaeal/bacterial primase domains (AEP) while the more C-terminal regions 3 and 4 showed similarities with DEAD and DEAH box helicases (37)). Analysis using hhpred (https://toolkit.tuebingen.mpg.de/tools/hhpred/) indicates that the N-terminal region (1-321) resembles the AEP Archaeo-Eukaryotic Prim-Pol Primase-Polymerase (43)(Fig. 4) while the C-terminal region (321-780) is similar to the ATP-dependent DNA helicase, UvsW, of bacteriophage T4. The authors find that targeted mutagenesis of the helicase domains “considerably reduces IEE activity” (37).

**Figure.3.**
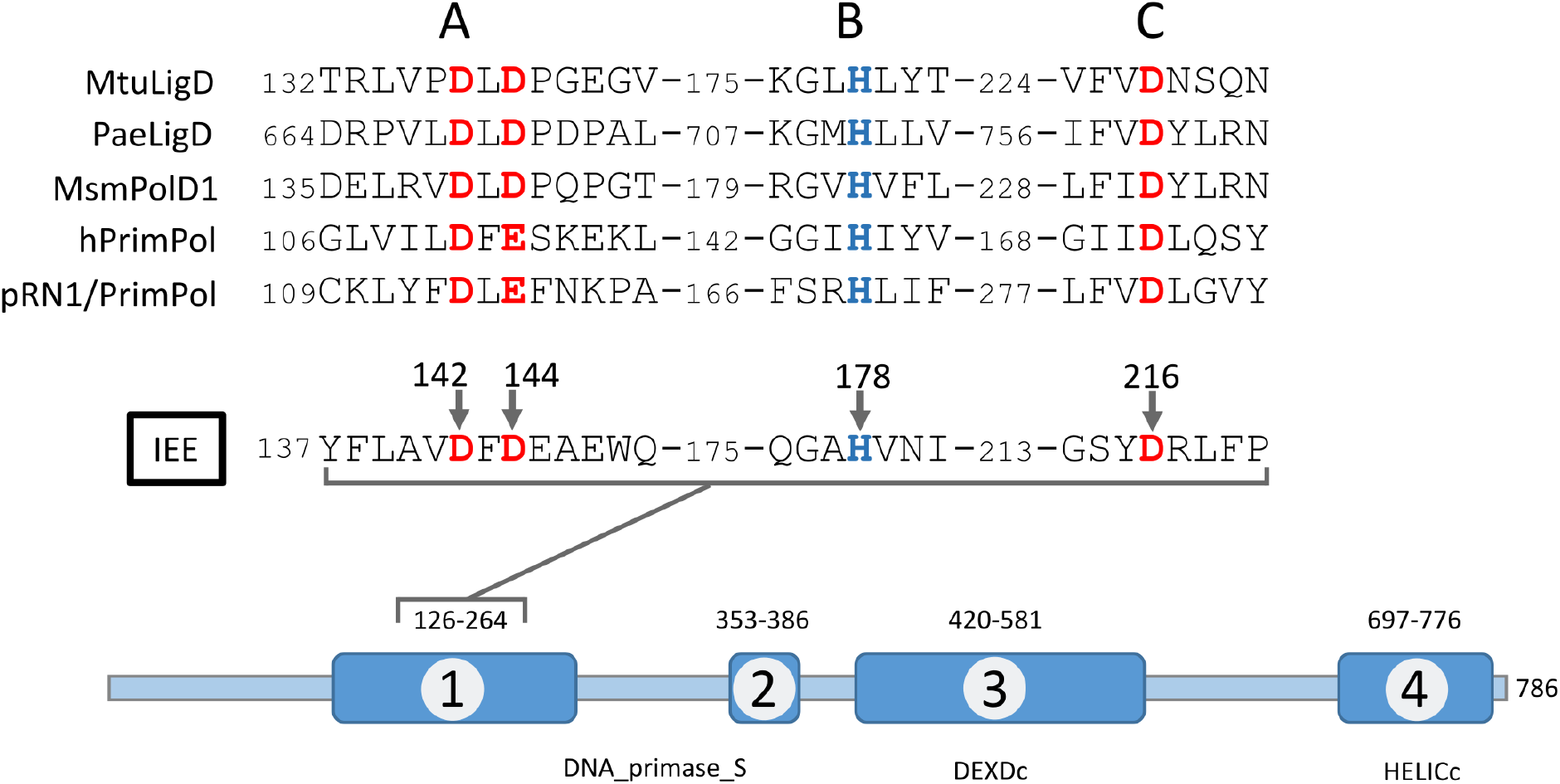
Organisation of the IEE protein. Schematic of the IS-excision enhancer protein is shown in blue. Alignment of a number of IEE proteins revealed 4 conserved regions whose position is indicated by the amino acid residues above the cartoon. Two of the more N-terminal regions (1 and 2) showed regions with similarities to eukaryotic/archaeal/bacterial primase domains (AEP), DNA primase_S (pfam01896) while the more C-terminal regions 3 and 4 show similarities DEAD and DEAH box helicases, DEXDc and HELICc .The four regions are indicated and sequence signatures are those identified by Kusumoto et al., 2011 PMID: 21224843. Three conserved regions, A, B and C, (Calvo et al 2023 PMID: 36715333) form the active site: Mg^2+^/Mn^2+^ ligands, D, are indicated in bold red letters; a conserved His which interacts with the incoming nucleotide is in blue. MtuLigD, *M. tuberculosis* NHEJ Ligase D; PaeLigD, P. *aeruginosa* NHEJ Ligase D; MsmPolD1, *M. smegmatis* PolD1; hPrimPol, human PrimPol ; pRN1/PrimPol, *Sulfolobus islandicus* plasmid pRN1 PrimPol. IEE Catalytic site residues are numbered D142, D144, H178, D 216.

**Figure.4.**
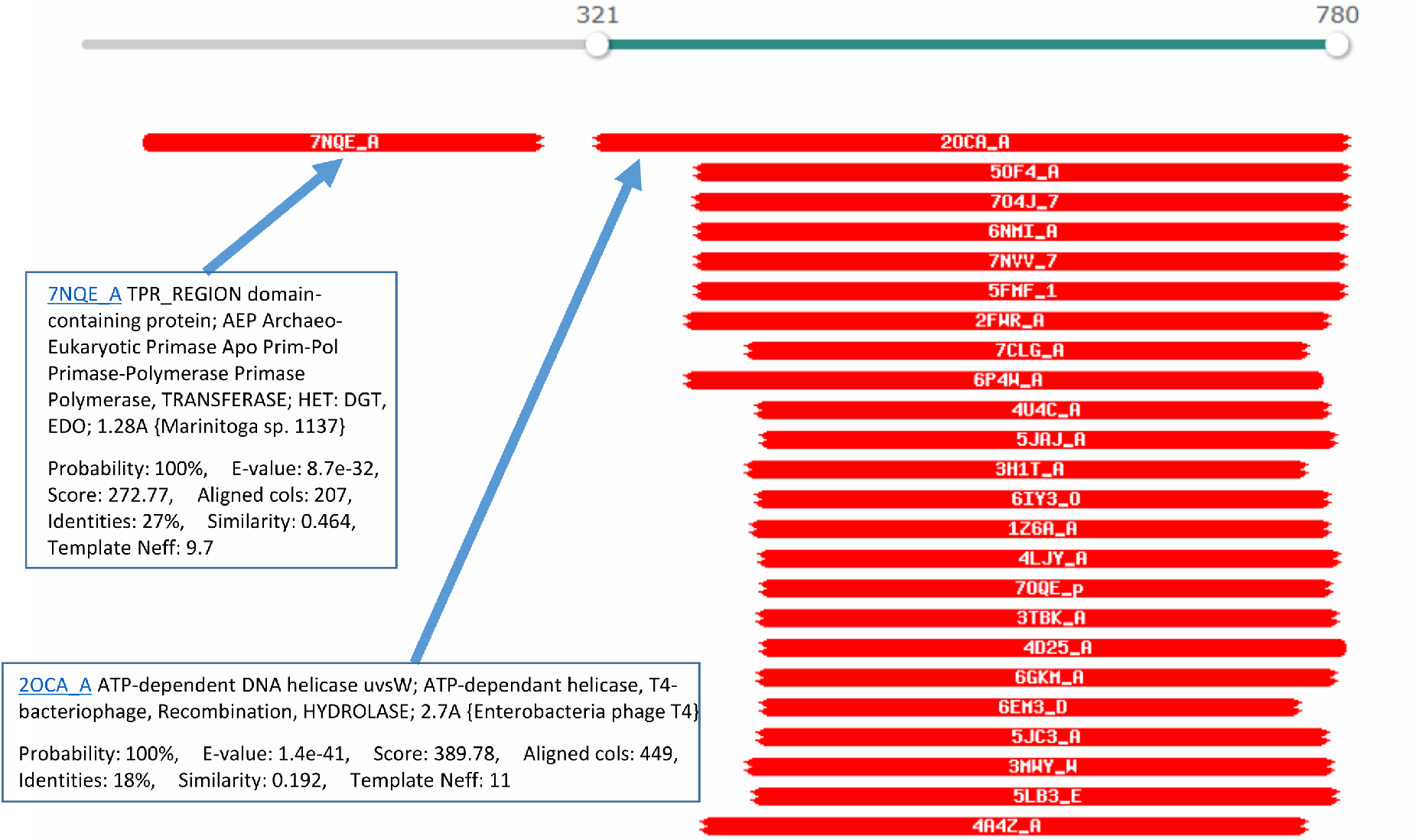
HhPRED Analysis of IEE. The N-terminal domain prior to amino acid residue 321 has the following predictive result: 7NQE_ATPR_REGION domain-containing protein; AEP Archaeo-Eukaryotic Primase Apo Prim-Pol Primase-polymerase, TRANSFERASE; HET: DGT, EDO; 1.28A (martinitogasp 1137) Probability: 100%, E-value: 8.7e-32, Score: 272.77, Aligned cols: 207, Identities: 27%, Similarity: 0.464, Template Neff: 9.7. The C-terminal domain following amino acid residue 321 has the following predictive result: 2OCA_AATP-dependent DNA helicase *uvsW*; ATP-dependant helicase, T4-bacteriophage, Recombination, HYDROLASE; 2.7A {Enterobacteriophage T4} Probability: 100%, E-value: 1.4e-41, Score: 389.78, aligned cols: 449, Identities: 18%, Similarity: 0.192, Template Neff: 11

### Biochemical Analysis of IEE Activities: Polymerase Activity

Indeed, as proposed from alignments with other members of this protein family (37,38) (Fig. 3), biochemical analyses using purified IEE demonstrated that the enzyme possesses DNA polymerase activity in the presence of Mg^2+^ (38) and can extend a 15 bp primer on a short 33bp template using dNTPs but not NTPs (Fig. 5A**i**). Mutation of two of the probable AEP catalytic residues (Fig. 3), D142A/D144A, eliminated this activity.

**Figure.5.**
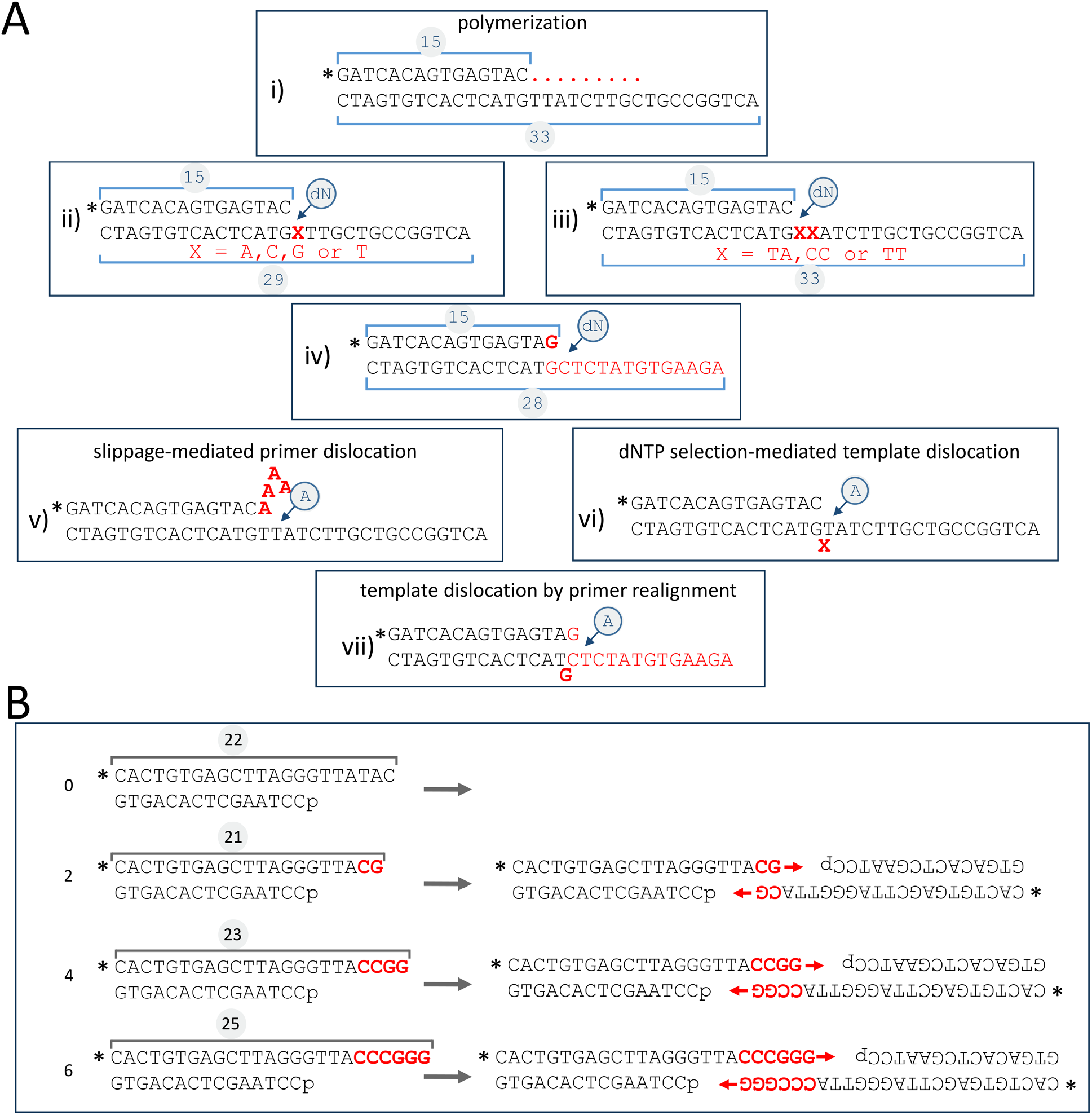
(A) Oligonucleotides Used in Assessing the Polymerization Activities of IEE. Taken from Calvo et al 2023 PMID: 36715333. All primers were 5’ end-labeled as shown by * i) polymerase extension reaction substrate. Extension shown as red dots. ii)And iii) incoming nucleotide selection substrates with a single X or double XX template nucleotide change. iv) substrate carrying a GG mismatch used in template realignment assay. v) slippage-mediated primer dislocation on templates such as i) or ii): addition of successive A residues (shown in red) on a T template dinucleotide when the reaction is provided with dA or on a C template dinucleotide when provided with dG. vi) dNTP selection-mediated template dislocation where IEE bypasses a template base (X in red). vii) template dislocation by primer realignment where the primer realigns with the template by extruding a template base (G in red). **(B) Oligonucleotides Used in Assessing IEE Microhomology Search Activities**. Taken from Calvo et al 2023 PMID: 36715333. All primers were 5’ end-labeled as shown by *. The left hand side shows the DNA substrates with the overlap region indicated in red. From top to bottom: 0, 2, 4 and six nucleotide overlaps. The right hand side shows the possible pairings using microhomology. The red arrows indicate primer extension used to assess the pairing. This generates a ladder of top strand oligonucleotides corresponding to the primer extension. Note that in some reactions DNA ligase was added and a small signal with the length expected of a joined single strand product was observed.

As might be expected from other examples of a number of endonucleases and nucleotidyl-transferases, the polymerization reaction was significantly higher in the presence of Mn^2+^ ions and the extension length was increased.

Experiments using various molar ratios of Mn^2+^ and Mg^2+^ led the authors speculate that the IEE AEP domain might preferentially use Mn^2+^ *in vivo* even in the presence of Mg^2+^. They argue that the active site of some AEPs is more flexible than that of replicative polymerases since it is configured to “*accommodate dislocations of the template and primer strands as well as to extend mismatched base pairs*” (38). These include the AEP domains of the Translesion Synthesis (TLS) class such as human PrimPol, and those of the Non Homologous End Joining (NHEJ) class such as bacterial LigD. They suggest that this flexibility allows preferential accommodation of Mn^2+^.

Preferential IEE use of Mn^2+^ suggested that it too could also allow extension with mismatched base pairs and promote DNA strand rearrangements during polymerization. This was demonstrated using a set of substrates carrying a single or double base changes in the template strand (Fig.5Aii and Aiii) while supplying individual defined nucleotides in the reaction. Although the enzyme showed a strong preference for incorporation of complementary nucleotides at the templating nucleotides (Fig.5Aii) during primer extension, a significant degree of misincorporation (∼16%) also occurred (38).

Interestingly, a major product resulted from insertion of two (identical) nucleotides on substrates in which the first and second templating nucleotides were different (Fig.5Aiii). In cases where these were the same (Fig.5Av), addition of the complementary nucleotide to the reaction resulted in an “expansion” by “reiterative” primer strand slippage to extend the primer by 5-7 nucleotides as shown in Fig.5Av. This was observed for both the TT and CC template pair where reiterative addition of A or G occurred respectively. IEE could also bypass the first template base in substrates (Fig.5Av) and bypass a mismatch (Fig.5Aiv) as schematized in Fig.5Avii.

The authors conclude that their results indicate that IEE is an “*error-prone DNA polymerase*” which can undergo slippage-mediated primer dislocation (Fig.5Av), dNTP selection-mediated template dislocation (Fig.5Avi) and template dislocation by primer realignment (Fig.5Avii), all resulting in DNA distortion. Dislocation of primer and template strands would, of course, facilitate the search for microhomologies.

### Microhomology-Mediated End-Joining

These properties raised the question of whether, like bacterial end-joining PolDom (eg. (44)), the primer/template dislocations catalyzed by the IEE AEP domain could promote Microhomology-Mediated End Joining, MMEJ). which might explain how the enzyme stimulates IS excision using its reduced nucleotide insertion fidelity (as shown in Fig.10).

Using appropriately 5’ resected double strand oligonucleotides (Fig.5B) with 0, 2, 4 and 6 nucleotide microhomologies at their 3’ ends (Fig.5B left). IEE-mediated MMEJ would create 3’ ends which should act as primers on 3’-synapsed DNA (Fig.5B right). A polymerization reaction including the four dNTPs revealed that IEE could specifically elongate a significant proportion of the 4 and 6 nucleotide substrates (28 and 46%) although no detectable elongation was observed with the 2 nt substrate under the reaction conditions used. IEE was also capable of limited synapsing of single strand substrates with 3’ terminal microhomologies (38).

### The C-terminal Helicase Domain

IEE exhibits DNA-dependent ATPase activity which is eliminated in an active site K451A mutant. However, unlike other ATP dependent helicases, it was not able to couple its ATPase activity with unwinding of double strand DNA with 3’ or 5’ unpaired tails (38). The role of the helicase domain remains unclear although a number of experiments comparing the activities of full length IEE and a C-terminal truncation which deletes the helicase domain from residue 289 (Fig.3) suggested that the C-terminal domain plays a role in stabilizing the interaction of the full length protein with its DNA substrates. The authors propose that it might even contribute to the removal of transposase from the transposition complex (38).

### Models for IEE activity

Using a non-selective PCR approach, IEE-enhanced excision was shown not only to increase precise excision but also other, more extensive, deletion events. It was hypothesized that these occurred as a consequence of the IS*3* family transposition mechanism in which initial cleavage of one IS end (Fig.6iA) is followed by its strand transfer close to the opposite end to form a bridged molecule containing a small flanking sequence from the vector plasmid at the junction (Fig.6iB) (6). The deletions were explained by the hypothesis that this initial strand transfer could be ‘sloppy’ and DNA sequences other than a second end could serve as a target (Fig.6i**i**).

**Figure.6.**
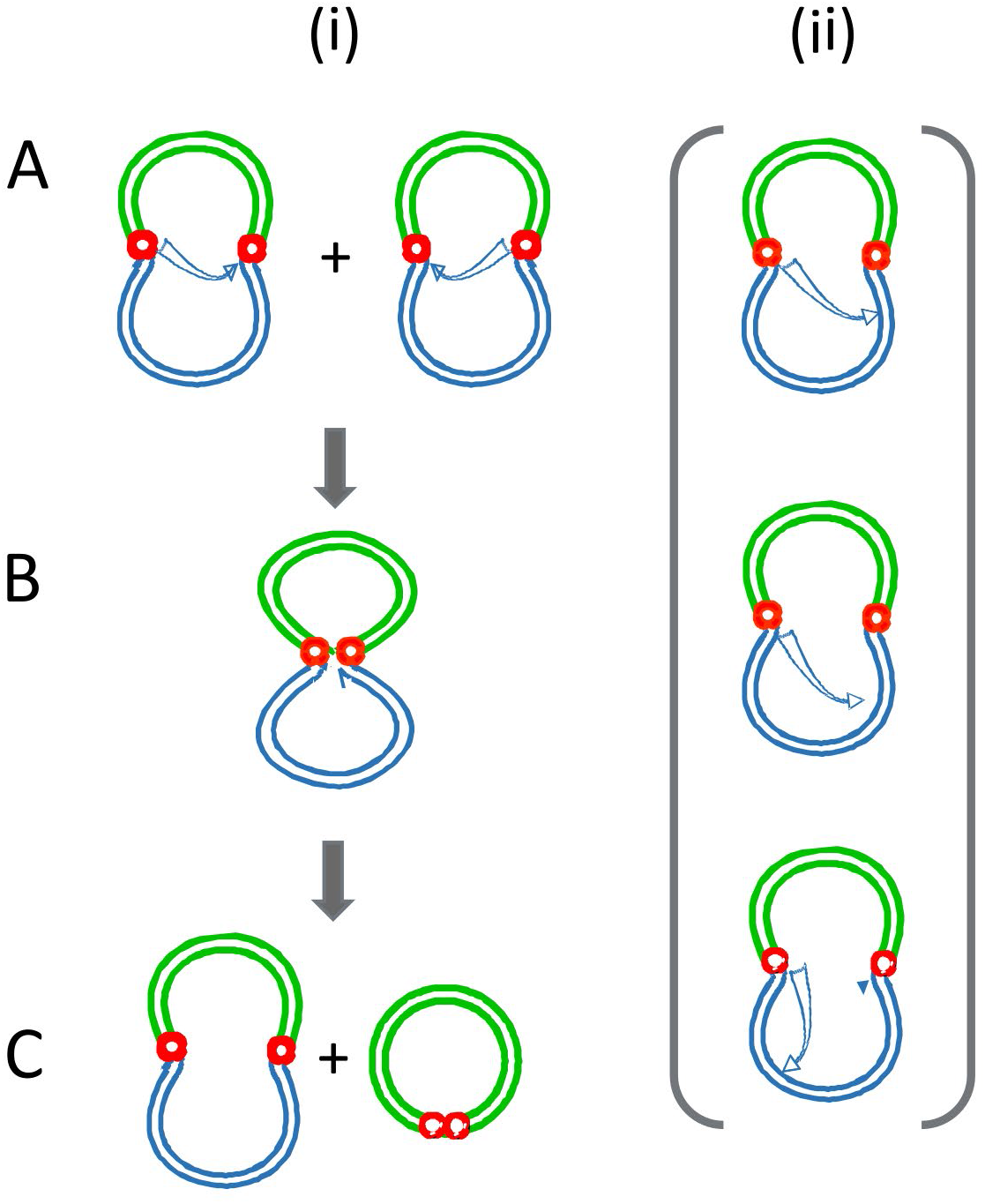
Model Proposed for generating different IS629 deletions. Redrawn and adapted from Chandler et al., 2015 PMID: 26350305 and Kusumoto et al 2011 PMID: 21224843. Transposon sequences are shown in green; transposon ends are shown as red circles and the neighboring host DNA as black lines. The arrows show attack by one IS end at the opposite end that occurs during copy-out-paste-in transposition. **i)** the normal productive pathway leading to formation of a circular IS intermediate and regenerating the original donor replicon. Note that no IS loss occurs. **ii)** “sloppy” attack at different positions in the donor replicon. Note that from what is known of the copy-out-paste-in transposition mechanism, this would not be expected to result in IS excision.

However, transposition of IS*3* family members occurs by a copy-out mechanism and is thought to regenerate the donor plasmid (Fig.6iC). Such deletions would not be consistent with this mechanism but would require an additional type of reaction to resolve the deleted donor.

Calvo and colleagues (38) provide a model for IEE activity based on their biochemical results in which they propose that IEE pairs two 5’ resected ends DNA ends in a reaction which is facilitated by the C-terminal helicase domain, allowing the AEP domain to accomplish a “*filling in*” polymerization reaction. The model invokes an intermediate which is thought to occur during a cut-out-paste-in transposition pathway which leaves a blunt ended double strand break in the donor DNA molecule (see (45)). Moreover, most IS generate short, direct target repeat sequences on insertion and would therefore provide 3’ terminal microhomologies upon 5’ resection.

However, in addition to IS*3* family members, Kusumoto et al., (37) show that IEE also stimulates excision of IS*1* and IS*30* family members (Fig. 8), both of which can transpose using a copy-out-paste-in mechanism (7,46). A number of the IS exhibit a measurable level of excision in the absence of the *iee* **gene**. Interestingly, it was proposed, based on identification of *in vivo* transposase-induced structures, that IS*1* can transpose using a number of alternative pathways (7): IS*1* shows a low basal level of excision which is further stimulated by IEE. It is possible that excision in the absence of *iee* occurs by a different mechanism such as replication fork slippage (47,48) as has been suggested to occur in the IS*4* family member, IS*10*. In this light, it is noteworthy that IS*4* itself shows a low level of *iee* independent excision which is not affected by *iee* (Fig.7). Additionally, IS*5*, IS*26* and IS*621*, none of which are known to use a copy-out-paste-in transposition mechanism, were not observed to excise even in the presence of *iee*.

**Figure.7.**
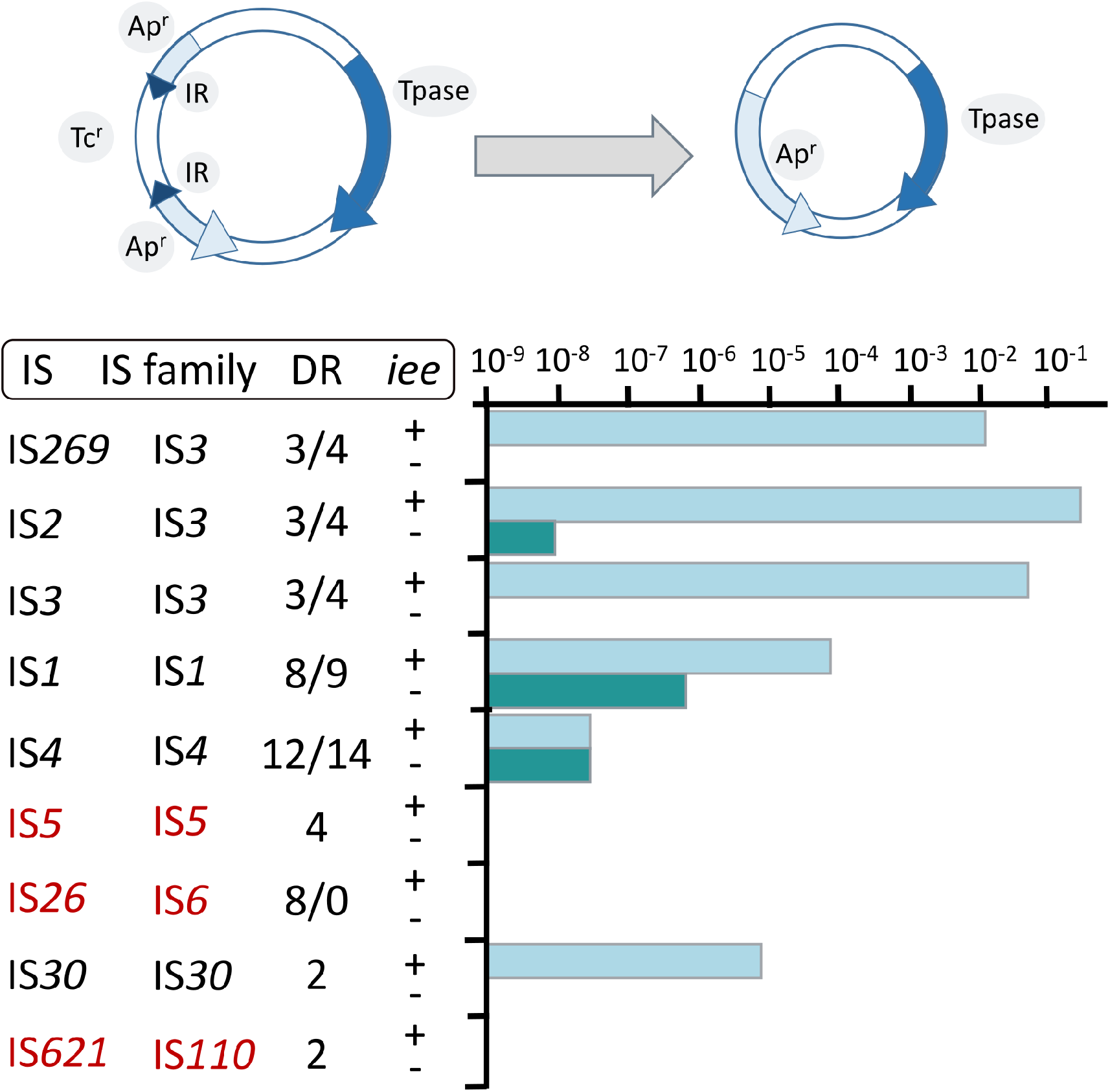
Relative IS Excision Frequencies. Redrawn and adapted from Kusumoto et al 2011 PMID: 21224843. A number of different IS were examined for IEE-stimulated excision in E.coli K-12 with and without a cloned *iee* copy. The assay consisted of a plasmid carrying an ampicillin resistance (Ap^r^) gene inactivated by insertion of a copy of the IS being tested whose transposase had been substituted for a tetracycline resistance gene (Tc^r^). The plasmid also included the transposase gene placed under control of an external promoter. Excision of the IS*629* derivative results in loss of Tc^r^ and appearance of Ap^r^.

**Figure.8.**
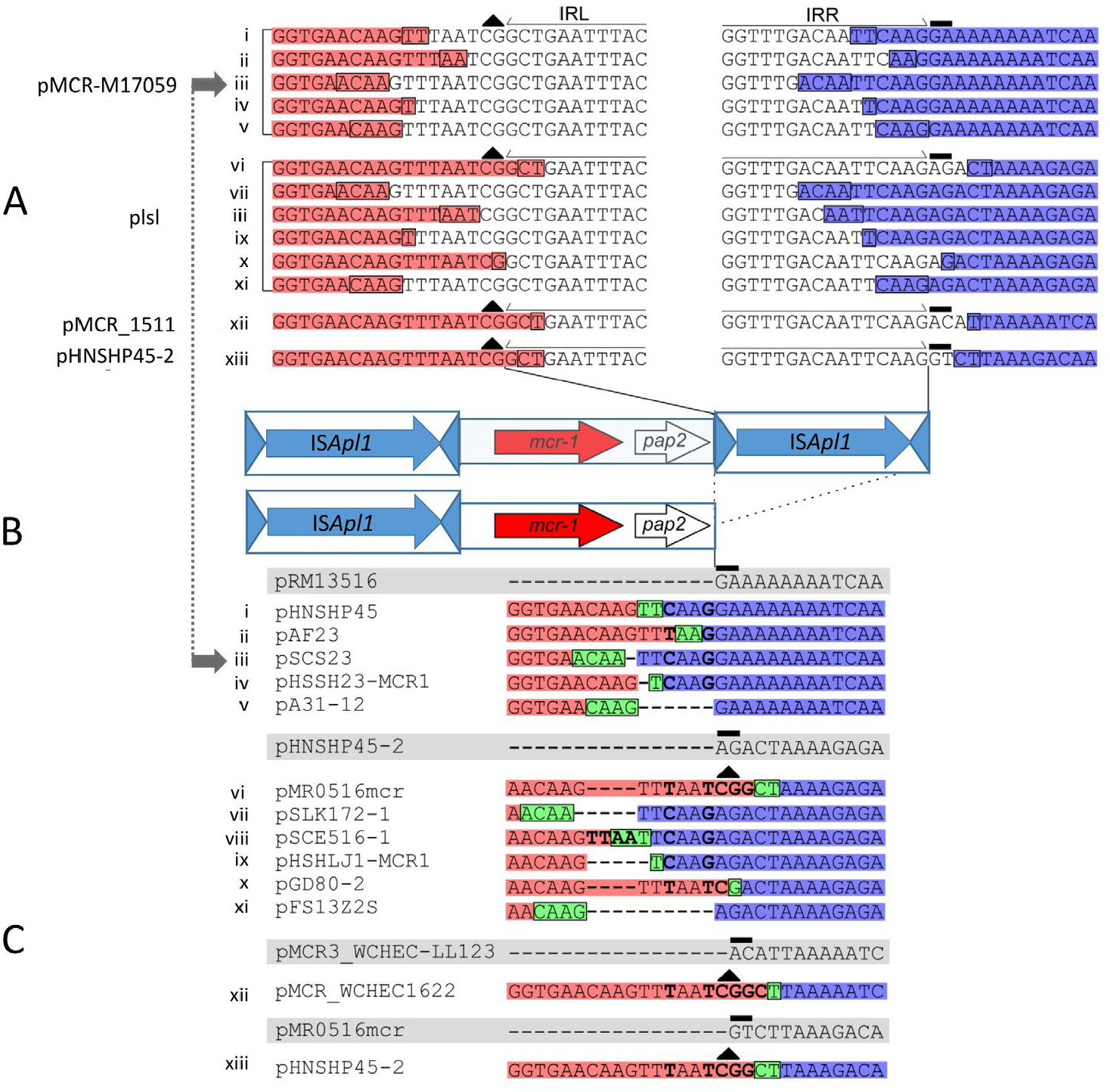
Alignment showing the decay of multiple different instances of Tn*6330*. The figure is redrawn and modified from Snesrud et al., 2018 PMID: 29440577. **(A) Sequence of four parental Tn*6330*-carrying plasmids**. The conserved, ancestral CG dinucleotide on the inside end (IE) of the downstream IS*Apl1* is indicated with a black triangle. The 2 bases at the end of the right-hand IS*Ap1* that are part of the DR generated by insertion of the entire Tn*6330* are over-scored in black. The half arrows above indicate the IS*Apl1* IRL and IRR sequences defining the ends of the downstream IS. The bases upstream and downstream that are retained after IS*Apl1* loss are highlighted in salmon and purple, respectively. The deletion joints upstream and downstream of the IS*Apl1* are boxed. The roman numerals on the left are correlated with the corresponding deletion product in C). **(B) Cartoon showing the structure of Tn*6330*** and, below, the generic structure of the product in which the downstream IS*Apl1* copy has been deleted. The IS is shown as a blue box. The triangles at each end represent the left (IRL) and right (IRR) terminal inverted repeats. The transposase open reading frame is shown by a blue horizontal arrow. *mcr-1* and *pap2* reading frames are shown as red and white horizontal arrows respectively. Thin black lines above, situate the downstream ISApl1 copy to the sequences in A) and the dotted lines below, to the deletion junction in C). **(C) Corresponding deletion products**. The sequences highlighted in grey show the sequence of plasmids with an empty site. The bases upstream and downstream of the deletion that are retained after IS*Apl1* loss are highlighted in red and blue, respectively, while the remaining copy of the deletion joint that is retained after the two ends are joined following IS*Apl1* excision is highlighted in green and encased in a black rectangle. The roman numerals on the left are correlated with the corresponding full length parent in A). The two horizontal blue arrows joined by a dotted blue line on the left indicate the parent and deletion product used in the example presented in Figure 10A.

Thus the common mechanistic property of those IS whose excision is stimulated by IEE is that they all form transposon circles and presumably use a copy-out pathway.

The simplest model for their excision mechanism is that excision products are generated, not during the first strand transfer step of copy-out-paste-in transposition as proposed (Fig.6iA), but during the second, replication step (copy-out) (Fig. 6iB), by slippage and realignment of the replication primer (see Fig. 10B). This had been proposed for the deletion of IS*30* family IS*Apl1* copies flanking the colistin resistance, *mcr*, gene in Tn*6330* (28).

### The mcr connection: a model for excision by replication-associated strand exchange

Colistin (polymixin E) is a last resort antibacterial that was used extensively in husbandry. Discovery of a transferable phosphethanolamine transferase conferring resistance to colistin in 2015 (49) was of such concern that it stimulated an immense effort to identify the resistance gene, *mcr*, in various bacterial sources worldwide. This quickly generated an extensive *mcr* sequence library in which it was noted that the gene was often, but not always, associated with an upstream or downstream copy of an IS*30* family sequence, IS*Apl1*. A number of examples carried two flanking IS copies, and the entire structure, which was proposed to be a compound transposon with characteristic 2bp direct target repeats (28), was called Tn*6330* (50) and subsequently confirmed to undergo transposition (29). Tn*6330* carries the *mcr1* gene together with a downstream open reading frame, *pap2*. Examination of a significant number of structures lacking the downstream IS revealed that the ⋃2.6 kb region including *mcr-1* and *pap2* was 99% identical; the non-identical nucleotides were concentrated at the 3’ end of *pap2*, the end that carries the downstream IS*Apl1* copy in Tn*6330* (see Fig.8). Moreover, AT rich the *pap2* gene in these cases was flanked by AT-rich regions. IS*Apl1* shows a strong preference for AT rich target sites, suggesting that an ancestral downstream IS*Apl1* copy had been deleted.

The number of available sequences was such that examples of closely related plasmid backbones could be identified which either had a complete Tn*6330* insertion, with “empty” sites (often in multiple examples) as well as cases in which examples also inserted into the same location in which one or the other flanking IS was absent.

Careful scrutiny of the sequences flanking *pap2* without an associated downstream IS, revealed small microhomologies which were thought to represent scars of deletions. Figure 8 shows 4 plasmids each containing Tn*6330* (Fig.8A). The salmon colored boxes on the left represent *pap2* sequences retained in the subsequent deletions and the purple boxes on the right represent the external flanking sequences retained (some of these intrude into the IS). Figure 8C shows individual “deletants” with similar backbones to those shown in Figure 8A. It was noted that, when aligned, the deletions have largely occurred between microhomologies of two to four base pairs. Similar results were obtained when the sequences upstream of mcr1 were examined. Interestingly, the length of the deleted segments (Fig.9) remains close to that of IS*Apl1*, 1070 bp.

**Figure.9.**
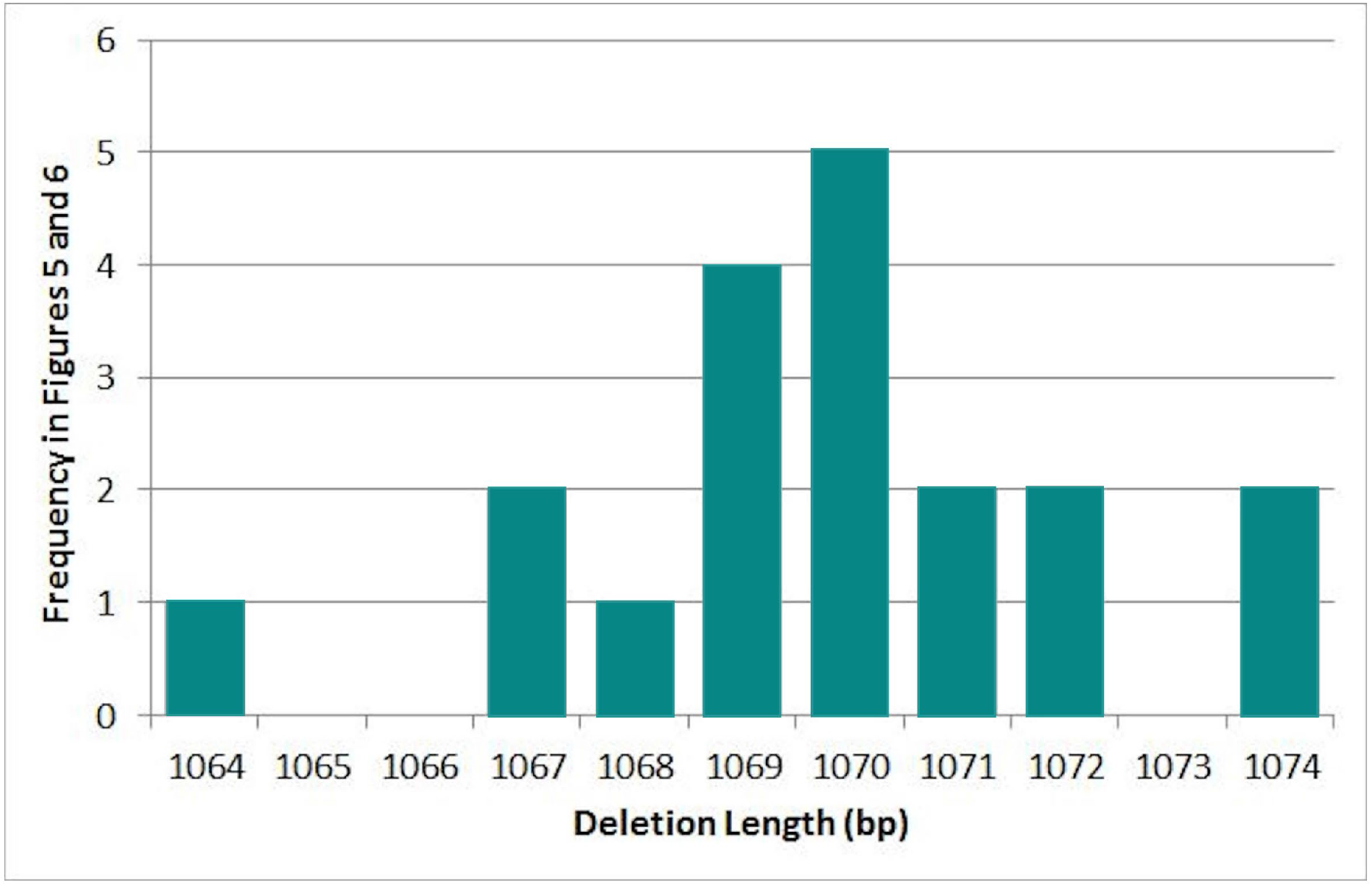
Distribution of different deletion sizes following loss of the downstream IS*Apl1* in 13 sequences in Figure 9 together with 6 obtained for deletion of the upstream ISpl1 copy. The average deletion size is 1,069.8 bp (Standard deviation: 2.4). From Snesrud et al., 2018 PMID: 29440577

A number of years earlier, Szabó et al. (51) had observed similar products with IS*30*. Additionally, when the IS*30* transposase gene was ablated, the deletion frequency was not only reduced by a factor of 10^3^ but the accompanying deletions were more complex, including large deletions or unidentified plasmid rearrangements.

The detailed observations obtained for IS*Apl1* led to the model shown in Figure 10B, which is illustrated with a specific example (iii in Fig.8) (Fig.10A).

**Figure.10.**
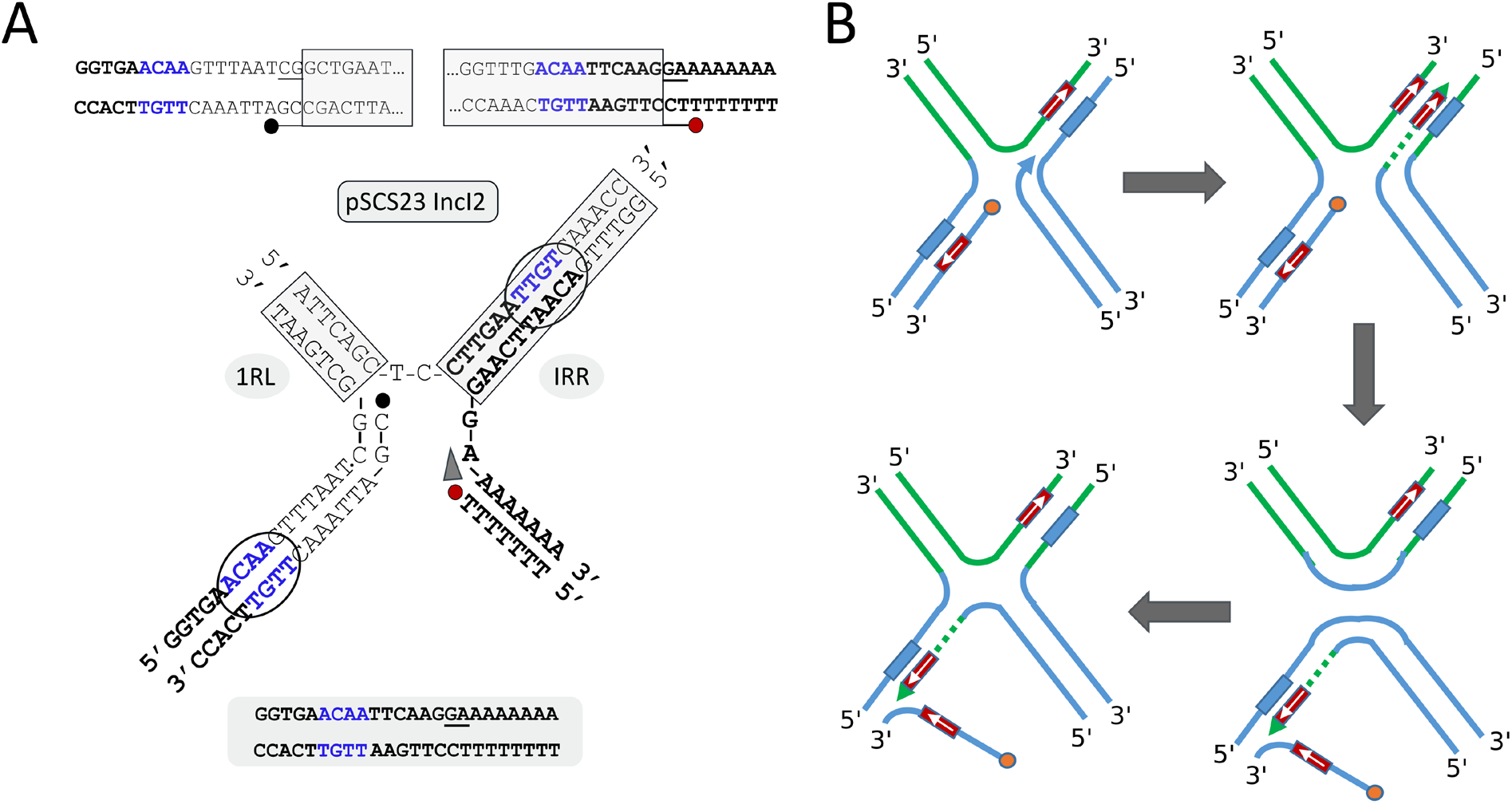
(A) A mechanism for IS*Apl1* deletion:The double-strand sequence of the IS ends in IncI2 plasmid pMCR-M17059 as an example. From Snesrud et al., 2018 PMID: 29440577. The panel presents the structure of the single-strand bridged molecule. IS ends are boxed in green. The 3’OH generated in the donor plasmid DNA is indicated by a red dot and the corresponding 5’ phosphate at the other IS end by a black dot. The blue arrow indicates the direction of transposition-associated replication. The deletion joint is shown in blue. The sequence remaining after deletion (bottom) representing plasmid pSCS23 is composed of the bold black characters together with one of the blue tetranucleotide sequences. **(B) A General Model showing how strand switching might lead to IS excision**. The model is based on data from analysis of loss of flanking IS*Apl1* elements from the colistin resistance compound transposon, Tn*6330*). Transposon DNA is shown in green, flanking DNA is shown in blue. The single strand bridged molecule is shown (from Figure 7iB) in which a short, complementary DNA sequence is represented by blue and magenta boxes with their relative orientation shown by a small blue arrow. **(i)** a 3’ primer from DNA neighboring the IS is indicated by the blue arrowhead, and the 5’ phosphate by the small orange circle. **(ii)** IS replication (copy-out) up to and including the duplicated sequence is indicated by a dotted green line. **(iii)** Strand switching of the primer strand to the duplicated sequence in the neighboring DNA is shown to occur with displacement of the complementary strand. This generates an intermediate which resembles a Holliday junction. **(iv)** Resolution of the Holliday junction resulting in deletion of most of the IS from the donor replicon. This structure can be resolved by replication.

In this model (Fig.10B), it is envisaged that a short complementary sequence occurring outside the IS is involved (Fig.10Bi) and that the DNA strand generated by transposition-associated replication (Fig.10Bii) switches to the complementary sequence (Fig.10Biii). This proposed structure is equivalent to a Holliday junction (see for example (52)), which could be resolved by RuvC (see for example (53)) (Fig.10Biv) resulting in the deletion of the intervening DNA.

This model is consistent with the results of IS*629* excision and might also be expected to involve a helicase and a primase able to locate microhomologies--exactly the biochemical properties demonstrated for the IEE PrimPol enzyme.

## Conclusion

In summary, the IEE protein plays an important role in excision of members of a number of IS families which all have in common the production of IS circles as transposition intermediates and probably all use the copy-out-paste-in transposition pathway. The excision is not only dependent on IEE but also requires an active transposase, indicating that it is associated with the transposition process itself. Not only do the biochemical properties of IEE include the ability to prime DNA synthesis and overcome potential obstacles due various lesions in the template DNA, they also include the capacity to recognize microhomologies. This suggests to us that IEE acts at the replication (copy-out) step in the transposition pathway, subsequent to initial 3’cleavage of on IS end and its transfer to the other. Specifically, based on sequence data obtained from the loss of the IS*30* family member, IS*Apl1*, which flanks the *mcr*-1 gene in the compound transposon Tn*6330*, we suggest that it allows a strand switch to suitable microhomologies in the neighboring donor DNA creating a Holliday junction and short circuiting the replication/transposition reaction. The IS could then be removed by Holliday junction resolution, a possibility that can be addressed experimentally.

Importantly, the activity of IEE in removing flanking IS serves to immobilize genes carried by compound transposons, explaining the presence of certain of these genes without associated IS copies in plasmids and chromosomes. It will be important to address this question experimentally using entire compound transposons such as Tn*6330* as well as others with flanking copy-out-paste-in IS and to compare the effects with compound transposons with different transposition pathways such as cut-and-paste.

## Bibliography

1. Siguier P, Gourbeyre E, Chandler M. Bacterial insertion sequences: their genomic impact and diversity. FEMS Microbiol Rev. 2014 Sep;38(5):865–891.

2. Siguier P, Perochon J, Lestrade L, Mahillon J, Chandler M. ISfinder: the reference centre for bacterial insertion sequences. Nucleic Acids Res. 2006 Jan 1;34(Database issue):D32–6.

3. Siguier P, Gourbeyre E, Varani A, Ton-Hoang B, Chandler M. Everyman’s guide to bacterial insertion sequences. Microbiol Spectr. 2015 Apr;3(2):MDNA3–0030.

4. Turlan C, Chandler M. Playing second fiddle: second-strand processing and liberation of transposable elements from donor DNA. Trends Microbiol. 2000 Jun;8(6):268–274.

5. Curcio MJ, Derbyshire KM. The outs and ins of transposition: from mu to kangaroo. Nat Rev Mol Cell Biol. 2003 Nov;4(11):865–877.

6. Chandler M, Fayet O, Rousseau P, Ton Hoang B, Duval-Valentin G. Copy-out-Paste-in Transposition of IS911: A Major Transposition Pathway. Microbiol Spectr. 2015 Aug;3(4).

7. Turlan C, Chandler M. IS1-mediated intramolecular rearrangements: formation of excised transposon circles and replicative deletions. EMBO J. 1995 Nov 1;14(21):5410–5421.

8. Sekine Y, Eisaki N, Kobayashi K, Ohtsubo E. Isolation and characterization of IS1 circles. Gene. 1997 Jun 3;191(2):183–190.

9. Polard P, Prère MF, Fayet O, Chandler M. Transposase-induced excision and circularization of the bacterial insertion sequence IS911. EMBO J. 1992 Dec;11(13):5079–5090.

10. Lewis LA, Grindley ND. Two abundant intramolecular transposition products, resulting from reactions initiated at a single end, suggest that IS2 transposes by an unconventional pathway. Mol Microbiol. 1997 Aug;25(3):517–529.

11. Arias-Palomo E, Berger JM. An Atypical AAA+ ATPase Assembly Controls Efficient Transposition through DNA Remodeling and Transposase Recruitment. Cell. 2015 Aug 13;162(4):860–871.

12. Kiss J, Olasz F. Formation and transposition of the covalently closed IS30 circle: the relation between tandem dimers and monomeric circles. Mol Microbiol. 1999 Oct;34(1):37–52.

13. Loessner I, Dietrich K, Dittrich D, Hacker J, Ziebuhr W. Transposase-dependent formation of circular IS256 derivatives in Staphylococcus epidermidis and Staphylococcus aureus. J Bacteriol. 2002 Sep;184(17):4709–4714.

14. Prudhomme M, Turlan C, Claverys JP, Chandler M. Diversity of Tn4001 transposition products: the flanking IS256 elements can form tandem dimers and IS circles. J Bacteriol. 2002 Jan;184(2):433–443.

15. Lauf U, Müller C, Herrmann H. Identification and characterisation of IS1383, a new insertion sequence isolated from Pseudomonas putida strain H. FEMS Microbiol Lett. 1999 Jan 15;170(2):407–412.

16. Higgins BP, Carpenter CD, Karls AC. Chromosomal context directs high-frequency precise excision of IS492 in Pseudoalteromonas atlantica. Proc Natl Acad Sci USA. 2007 Feb 6;104(6):1901–1906.

17. Prosseda G, Latella MC, Casalino M, Nicoletti M, Michienzi S, Colonna B. Plasticity of the P junc promoter of ISEc11, a new insertion sequence of the IS1111 family. J Bacteriol. 2006 Jul;188(13):4681–4689.

18. Henderson DJ, Lydiate DJ, Hopwood DA. Structural and functional analysis of the mini-circle, a transposable element of Streptomyces coelicolor A3(2). Mol Microbiol. 1989 Oct;3(10):1307–1318.

19. Smokvina T, Henderson DJ, Melton RE, Brolle DF, Kieser T, Hopwood DA. Transposition of IS117, the 2.5 kb Streptomyces coelicolor A3(2) “minicircle”: roles of open reading frames and origin of tandem insertions. Mol Microbiol. 1994 May;12(3):459–468.

20. Guérillot R, Siguier P, Gourbeyre E, Chandler M, Glaser P. The diversity of prokaryotic DDE transposases of the mutator superfamily, insertion specificity, and association with conjugation machineries. Genome Biol Evol. 2014 Feb;6(2):260–272.

21. Christie-Oleza JA, Nogales B, Lalucat J, Bosch R. TnpR encoded by an ISPpu12 isoform regulates transposition of two different ISL3-like insertion sequences in Pseudomonas stutzeri after conjugative interaction. J Bacteriol. 2010 Mar;192(5):1423–1432.

22. Vertes AA, Asai Y, Inui M, Kobayashi M, Yukawa H. The corynebacterial insertion sequence IS31831 promotes the formation of an excised transposon fragment. Biotechnol Lett. 1995 Nov;17(11):1143–1148.

23. Hudson CM, Bent ZW, Meagher RJ, Williams KP. Resistance determinants and mobile genetic elements of an NDM-1-encoding Klebsiella pneumoniae strain. PLoS One. 2014 Jun 6;9(6):e99209.

24. Nielsen TK, Rasmussen M, Demanèche S, Cecillon S, Vogel TM, Hansen LH. Evolution of Sphingomonad Gene Clusters Related to Pesticide Catabolism Revealed by Genome Sequence and Mobilomics of Sphingobium herbicidovorans MH. Genome Biol Evol. 2017 Sep 1;9(9):2477–2490.

25. Kusumoto M, Suzuki R, Nishiya Y, Okitsu T, Oka M. Host-dependent activation of IS1203v excision in Shiga toxin-producing Escherichia coli. J Biosci Bioeng. 2004;97(6):406–411.

26. Snesrud E, Ong AC, Corey B, Kwak YI, Clifford R, Gleeson T, et al. Analysis of Serial Isolates of mcr-1-Positive Escherichia coli Reveals a Highly Active ISApl1 Transposon. Antimicrob Agents Chemother. 2017 May;61(5).

27. Snesrud E, McGann P, Chandler M. The Birth and Demise of the ISApl1-mcr-1-ISApl1 Composite Transposon: the Vehicle for Transferable Colistin Resistance. MBio. 2018 Feb 13;9(1).

28. Snesrud E, He S, Chandler M, Dekker JP, Hickman AB, McGann P, et al. A Model for Transposition of the Colistin Resistance Gene mcr-1 by ISApl1. Antimicrob Agents Chemother. 2016 Oct 21;60(11):6973–6976.

29. Poirel L, Kieffer N, Nordmann P. In Vitro Study of ISApl1-Mediated Mobilization of the Colistin Resistance Gene mcr-1. Antimicrob Agents Chemother. 2017 Jul;61(7).

30. Kusumoto M, Nishiya Y, Kawamura Y. Reactivation of insertionally inactivated Shiga toxin 2 genes of Escherichia coli O157:H7 caused by nonreplicative transposition of the insertion sequence. Appl Environ Microbiol. 2000 Mar;66(3):1133–1138.

31. Hayashi T, Makino K, Ohnishi M, Kurokawa K, Ishii K, Yokoyama K, et al. Complete genome sequence of enterohemorrhagic Escherichia coli O157:H7 and genomic comparison with a laboratory strain K-12. DNA Res. 2001 Feb 28;8(1):11–22.

32. Makino K, Ishii K, Yasunaga T, Hattori M, Yokoyama K, Yutsudo CH, et al. Complete nucleotide sequences of 93-kb and 3.3-kb plasmids of an enterohemorrhagic Escherichia coli O157:H7 derived from Sakai outbreak. DNA Res. 1998 Feb 28;5(1):1–9.

33. Polard P, Prère MF, Chandler M, Fayet O. Programmed translational frameshifting and initiation at an AUU codon in gene expression of bacterial insertion sequence IS911. J Mol Biol. 1991 Dec 5;222(3):465–477.

34. Sekine Y, Nagasawa H, Ohtsubo E. Identification of the site of translational frameshifting required for production of the transposase encoded by insertion sequence IS 1. Mol Gen Genet. 1992 Nov;235(2-3):317–324.

35. Sekine Y, Ohtsubo E. DNA sequences required for translational frameshifting in production of the transposase encoded by IS1. Mol Gen Genet. 1992 Nov;235(2-3):325–332.

36. Chandler M, Fayet O. Translational frameshifting in the control of transposition in bacteria. Mol Microbiol. 1993 Feb;7(4):497–503.

37. Kusumoto M, Ooka T, Nishiya Y, Ogura Y, Saito T, Sekine Y, et al. Insertion sequence-excision enhancer removes transposable elements from bacterial genomes and induces various genomic deletions. Nat Commun. 2011 Jan 11;2:152.

38. Calvo PA, Mateo-Cáceres V, Díaz-Arco S, Redrejo-Rodríguez M, de Vega M. The enterohemorrhagic Escherichia coli insertion sequence-excision enhancer protein is a DNA polymerase with microhomology-mediated end-joining activity. Nucleic Acids Res. 2023 Jan 30;

39. Kusumoto M, Fukamizu D, Ogura Y, Yoshida E, Yamamoto F, Iwata T, et al. Lineage-specific distribution of insertion sequence excision enhancer in enterotoxigenic Escherichia coli isolated from swine. Appl Environ Microbiol. 2014 Feb;80(4):1394–1402.

40. Lundblad V, Kleckner N. Mutants of Escherichia coli K12 which affect excision of transposon Tn10. Basic Life Sci. 1982;20:245–258.

41. Lundblad V, Taylor AF, Smith GR, Kleckner N. Unusual alleles of recB and recC stimulate excision of inverted repeat transposons Tn10 and Tn5. Proc Natl Acad Sci USA. 1984 Feb;81(3):824–828.

42. Lundblad V, Kleckner N. Mismatch repair mutations of Escherichia coli K12 enhance transposon excision. Genetics. 1985 Jan;109(1):3–19.

43. Rudd SG, Bianchi J, Doherty AJ. PrimPol-A new polymerase on the block. Mol Cell Oncol. 2014 Jun;1(2):e960754.

44. Doherty AJ, Jackson SP, Weller GR. Identification of bacterial homologues of the Ku DNA repair proteins. FEBS Lett. 2001 Jul 6;500(3):186–188.

45. Benjamin HW, Kleckner N. Excision of Tn10 from the donor site during transposition occurs by flush double-strand cleavages at the transposon termini. Proc Natl Acad Sci USA. 1992 May 15;89(10):4648–4652.

46. Szabó M, Kiss J, Nagy Z, Chandler M, Olasz F. Sub-terminal sequences modulating IS30 transposition in vivo and in vitro. J Mol Biol. 2008 Jan 11;375(2):337–352.

47. Ross DG, Swan J, Kleckner N. Nearly precise excision: a new type of DNA alteration associated with the translocatable element Tn10. Cell. 1979 Apr;16(4):733–738.

48. Foster TJ, Lundblad V, Hanley-Way S, Halling SM, Kleckner N. Three’ ‘ Tn10-Associated Excision Events: Relationship to Transposition and Role of Direct and Inverted Repeats . Cell. 1981;23:215–227.

49. Liu Y-Y, Wang Y, Walsh TR, Yi L-X, Zhang R, Spencer J, et al. Emergence of plasmid-mediated colistin resistance mechanism MCR-1 in animals and human beings in China: a microbiological and molecular biological study. Lancet Infect Dis. 2016 Feb;16(2):161–168.

50. Li R, Xie M, Zhang J, Yang Z, Liu L, Liu X, et al. Genetic characterization of mcr-1-bearing plasmids to depict molecular mechanisms underlying dissemination of the colistin resistance determinant. J Antimicrob Chemother. 2017;72(2):393–401.

51. Szabó M, Kiss J, Kótány G, Olasz F. Importance of illegitimate recombination and transposition in IS30-associated excision events. Plasmid. 1999 Nov;42(3):192–209.

52. Lilley DMJ. Holliday junction-resolving enzymes-structures and mechanisms. FEBS Lett. 2017 Jan 1;591(8):1073–1082.

53. Kuzminov A. RuvA, RuvB and RuvC proteins: cleaning-up after recombinational repairs in E. coli. Bioessays. 1993 May;15(5):355–358.

54. Nakamura K, Seto K, Isobe J, Taniguchi I, Gotoh Y, Hayashi T. Insertion Sequence (IS)-Excision Enhancer (IEE)-Mediated IS Excision from the lacZ Gene Restores the Lactose Utilization Defect of Shiga Toxin-Producing Escherichia coli O121:H19 Strains and Is Responsible for Their Delayed Lactose Utilization Phenotype. Appl Environ Microbiol. 2022 Aug 23;88(16):e0076022.

55. Gill A, McMahon T, Dussault F, Jinneman K, Lindsey R, Martin H, et al. Delayed lactose utilization among Shiga toxin-producing Escherichia coli of serogroup O121. Food Microbiol. 2022 Apr;102:103903.

